# Internalization of erythrocyte acylpeptide hydrolase is required for asexual replication of *Plasmodium falciparum*

**DOI:** 10.1101/533570

**Authors:** Rubayet Elahi, Christie Dapper, Michael Klemba

## Abstract

The human malaria parasite *Plasmodium falciparum* causes disease as it replicates within the host’s erythrocytes. We have found that an erythrocyte serine hydrolase, acylpeptide hydrolase (APEH), accumulates within developing asexual parasites. Internalization of APEH was associated with a proteolytic event that reduced the size of the catalytic polypeptide from 80 to 55 kDa, which suggests that the enzyme resides in the food vacuole. A triazole urea APEH inhibitor, termed AA74-1, was employed to characterize the role of parasite-internalized APEH. *In vitro*, AA74-1 was a potent and highly selective inhibitor of both host erythrocyte and parasite-internalized APEH. When added to cultures of parasite-infected erythrocytes, AA74-1 was a relatively poor inhibitor of replication over one asexual replication cycle; however, its potency increased dramatically after a second cycle. This enhancement of potency was not abrogated by the addition of exogenous isopentenyl pyrophosphate, which distinguishes it from the well-characterized “delayed death” phenomenon that is observed with inhibitors that target the parasite apicoplast. Analysis of inhibition by AA74-1 *in vivo* revealed that a concentration of 100 nM was sufficient to quantitatively inhibit erythrocyte APEH. In contrast, the parasite-internalized APEH pool was inefficiently inhibited at concentrations up to 100-fold higher. Together, these findings provide evidence for an essential catalytic role for parasite-internalized APEH and suggest a model for AA74-1 growth inhibition whereby depletion of parasite APEH activity requires the internalization of inactive host cell APEH over two replication cycles.

**IMPORTANCE:** Nearly half a million deaths were attributed to malaria in 2017. Protozoan parasites of the genus *Plasmodium* cause disease in humans while replicating asexually within the host’s erythrocytes, with *P. falciparum* responsible for most of the mortality. Understanding how *Plasmodium* spp. has adapted to its unique host erythrocyte environment is important for developing malaria control strategies. Here, we demonstrate that *P. falciparum* co-opts a host erythrocyte serine hydrolase termed acylpeptide hydrolase. By showing that the parasite requires acylpeptide hydrolase activity for replication, we expand our knowledge of host cell factors that contribute to robust parasite growth.

## INTRODUCTION

In 2017, an estimated US $3.1 billion was spent on malaria control worldwide. Despite this expenditure, around half a million deaths due to malaria were reported that year (1). *Plasmodium falciparum*, one of the five species that cause human malaria, accounts for the vast majority of these deaths (1). While still unacceptably large, the latest mortality figure represents a substantial improvement on the malaria situation of fifteen years ago, which is due in part to the implementation of artemisinin combination therapy (2). Recent reports of reduced efficacy in Southeast Asia have raised concerns that parasites are evolving resistance (or tolerance) to artemisinins and their partner drugs (3, 4). The discovery and validation of new anti-malarial targets is therefore a critical component of a robust anti-malarial pipeline, which is needed to safeguard recent advances and to devise strategies for eradication.

Enzymes of the serine hydrolase superfamily encompass a highly diverse range of catalytic activities and have garnered much attention for their roles in many critical metabolic processes in humans (5, 6). Based on annotated sequence homologies, the *P. falciparum* genome encodes over 40 putative members of the serine hydrolase superfamily (7), most of which have not been functionally characterized. Exploration of the roles of uncharacterized serine hydrolases will lead to new insights into essential aspects of parasite metabolism and possibly to new chemotherapeutic targets.

Serine hydrolase-directed activity-based probes (ABPs) have emerged as powerful tools for the functional annotation of serine hydrolases in complex proteomes (5, 8). By enabling competitive activity-based protein profiling (ABPP), ABPs have greatly accelerated the discovery of inhibitors that are highly specific for individual serine hydrolases (5). ABPs containing a fluorophosphonate (FP) warhead provide broad coverage of the serine hydrolase superfamily with negligible off-target activity (9, 10). Reaction of the FP warhead with the active site serine forms a stable covalent adduct. ABPs containing a fluorescent reporter enable a direct quantitative readout of the levels of active serine hydrolases (10).

We have employed a fluorescent FP probe in conjunction with well-characterized serine hydrolase inhibitors to profile the serine hydrolase activities of asexual intraerythrocytic *P. falciparum*. In the course of these studies, we made the surprising discovery that a human host erythrocyte serine hydrolase, acylpeptide hydrolase (APEH, EC: 3.4.19.1; also referred to as acylamino acid releasing enzyme and acylaminoacyl-peptidase) is one of the most abundant serine hydrolases in the developing asexual parasite. APEH is a member of the prolyl oligopeptidase (POP) family of serine peptidases (clan SC, family S9C). In mammals, APEH is ubiquitously expressed (11) and it has been purified from human erythrocytes as a homotetramer (12, 13). APEH was initially identified as an exopeptidase that catalyzes the hydrolysis of N-terminally acylated amino acids from peptides, yielding an acylamino acid and a shortened peptide with a free N-terminus (14, 15). Acetylated and formylated peptides are good substrates for APEH (16, 17). There have also been reports of APEH endopeptidase activity against oxidized proteins (18) and amyloidogenic Aβ peptide (19).

The physiological roles of APEH in mammalian cells are not completely understood. On the basis of the exopeptidase activity of APEH noted above, it has long been hypothesized that APEH participates in the maturation of proteins through the removal of acetylated N-terminal residues (20). Treatment of mouse T cells with a potent and highly selective inhibitor of APEH, termed AA74-1, affected the acetylation status of 25 proteins, lending support for this hypothesis (21). There is some evidence that APEH influences activity of the proteasome (22, 23); however, the mechanistic details of this interaction remain to be elucidated.

Here, we have employed the fluorescent activity-based serine hydrolase probe TAMRA-fluorophosphonate (TAMRA-FP), the covalent triazole urea APEH inhibitor AA74-1, and anti-APEH antibodies to explore the properties and physiological role of the parasite-internalized enzyme.

## RESULTS

### Identification of human APEH in saponin-isolated *P. falciparum*

As a first step towards a proteome-wide functional annotation of serine hydrolase activities in asexual *P. falciparum*, we compared the TAMRA-FP labeling profiles of crude lysates of uninfected erythrocytes and of saponin-isolated parasites (Fig. 1A). Saponin selectively permeabilizes the erythrocyte plasma membrane and the parasitophorous vacuole (PV) membrane of parasite-infected red blood cells; thus, saponin-treated parasites lack soluble erythrocyte and PV proteins (24). Unsurprisingly, there is little overlap between the two profiles, which is consistent with an organism-specific pattern of serine hydrolase expression.

**Figure 1:**
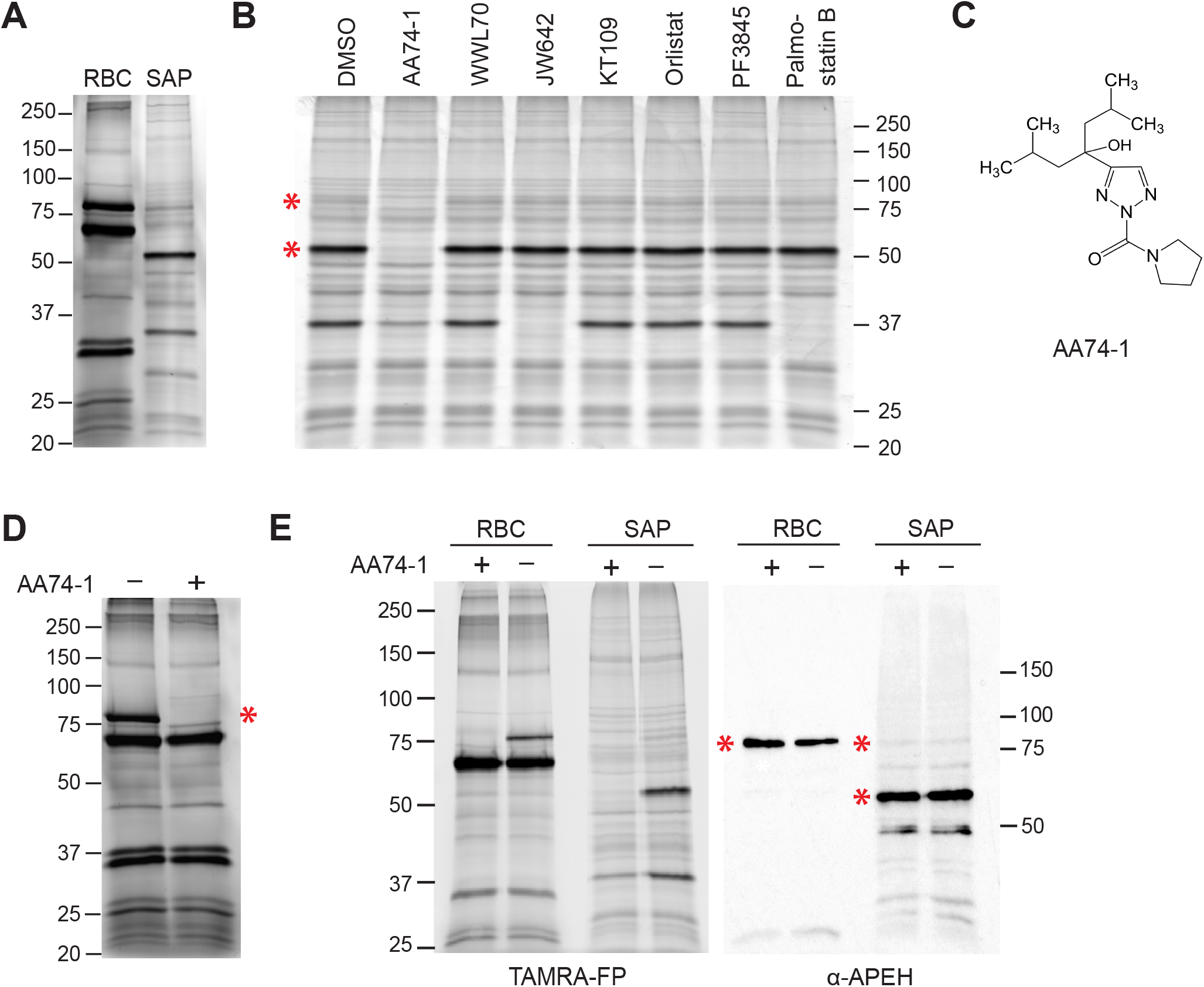
Identification of human APEH in saponin-isolated *P. falciparum*. (A) TAM-RA-FP labeling of serine hydrolases in crude lysates of uninfected human erythrocytes (RBC) and saponin-isolated parasites (SAP). (B) Competitive ABPP using a panel of serine hydrolase inhibitors (1 µM) that target various classes of serine hydrolase (see Table S1) or vehicle (1% DMSO) control. The 80 and 55 kDa species that react quantitatively with AA74-1 are indicated with red asterisks. (C) Structure of the APEH-selective inhibitor AA74-1. (D) Effect of AA74-1 (1 µM) on TAMRA-FP labeling of serine hydrolases in a crude lysate of uninfected erythrocytes. The red asterisk indicates an 80 kDa species that is inhibited by AA74-1. (E) Immunodetection of human APEH. Left panel: TAMRA-FP profile of serine hydrolases in lysates of uninfected erythrocytes (RBC) and saponin-isolated parasites (SAP) with and without 1 µM AA74-1. Right panel: anti-APEH immunoblot following transfer of proteins to nitrocellulose. Red asterisks indicate AA74-1-inhibited species. Bands below 50 kDa in SAP lanes may represent APEH degradation products. Sizes of molecular markers are indicated in kDa.

To gain insight into the functionality of *P. falciparum* serine hydrolases, competitive activity-based probe profiling (referred to as competitive ABPP) was conducted. Crude lysates of saponin-isolated parasites were incubated with covalent inhibitors of diverse human serine hydrolases prior or vehicle (1% DMSO) prior to TAMRA-FP labeling (Fig. 1B; structures of inhibitors and their known targets are provided in Table S1). We were intrigued to find that AA74-1, a triazole urea inhibitor (Fig. 1C) that is highly selective for human APEH (21), completely blocked TAMRA-FP labeling of a major ∼55 kDa species and a minor ∼80 kDa species in parasite lysate (Fig. 1B). In contrast, lipase or fatty acid amide hydrolase inhibitors did not compete with labeling of the 55 or 80 kDa species (Fig. 1B). Treatment of erythrocyte lysate with the same inhibitor panel revealed that AA74-1, but not lipase or fatty acid amide hydrolase inhibitors, blocked TAMRA-FP labeling of an ∼80 kDa species in a highly selective manner (Figs. 1D, S1). The estimated molecular mass of this species is consistent with a predicted molecular mass of 81.2 kDa for human APEH (25).

The above findings suggest two possible interpretations: i) *P. falciparum* expresses an endogenous protein with activity similar to that of human APEH, or ii) the parasite internalizes the erythrocyte enzyme. To distinguish between these possibilities, we asked whether parasite-internalized APEH is recognized by an affinity purified anti-human APEH antibody (Fig. 1E). Parasite and erythrocyte lysates were first analyzed by competitive ABPP with and without AA74-1 to identify APEH (Fig. 1E). After in-gel fluorescence scanning of TAMRA-FP-labeled species, the proteins were transferred to nitrocellulose and APEH was detected by immunoblotting. The 80 kDa species in erythrocyte lysate and the major 55 kDa and minor 80 kDa species in parasite lysate were recognized by the antibody, with the relative abundance of the species on the membrane comparable to that observed in the TAMRA-FP scan (Fig. 1E). These findings strongly suggest that *P. falciparum* internalizes APEH, which then appears to undergo a proteolytic event to reduce the size of the active site-containing segment from 80 to 55 kDa. The minor ∼80 kDa species in parasite lysate that is inhibited by AA74-1 likely represents full-length, uncleaved APEH. Hereafter, these two species will be collectively referred to as “parasite-internalized APEH”.

### Validation of AA74-1 as a potent and selective inhibitor of parasite-internalized APEH

Before using AA74-1 to probe the importance of internalized APEH for intraerythrocytic parasite development, we evaluated its potency and selectivity *in vitro*. The 50% inhibitory concentration (IC_50_) values for AA74-1 inhibition of erythrocyte and parasite-internalized APEH were determined by competitive ABPP (Fig. 2A, B). Mean IC_50_ values from three independent replicates were 7.9 ± 1.8 nM for the parasite 55 kDa species and 7.4 ± 2.4 nM for the erythrocyte 80 kDa species, which are not significantly different (two-tailed Student’s *t*-test, *p*-value = 0.78). These values are very close to the 11 nM IC_50_ value reported for AA74-1 inhibition of APEH in a human cell line using a similar competitive ABPP assay (21). The parasite 80 kDa species appeared to have a comparable IC_50_ value (Fig. 2A), but its lower abundance made it difficult to reliably quantify this species.

**Figure 2:**
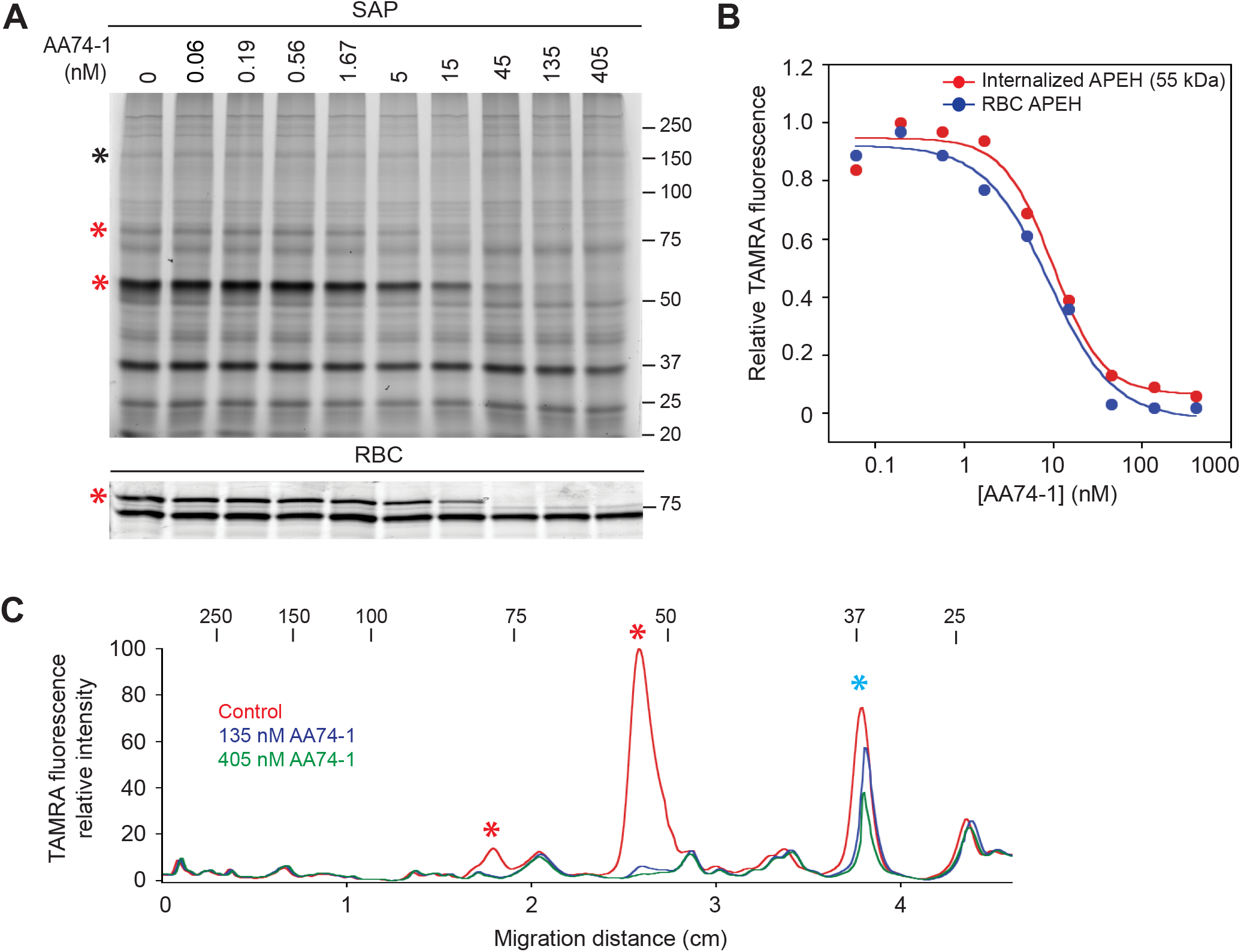
*In vitro* potency and selectivity of AA74-1 for parasite-internalized APEH. (A) Competitive ABPP with AA74-1 over a concentration range of 0.06 – 405 nM. Upper panel: Saponin-isolated parasites (SAP). The 55 and 80 kDa APEH species are indicated with red asterisks. The species used for APEH peak volume normalization (see Materials and Methods) is indicated with a black asterisk. Lower panel: Uninfected erythrocytes (RBC). APEH is indicated with a red asterisk. See Fig. S2 for the full gel image and the species used for peak volume normalization. (B) Plot of normalized APEH peak volume, expressed as a fraction of the control (“0 nM”, 1% DMSO), vs. AA74-1 concentration. Data are from the gel image in (A), which represents one of three biological replicates. Points were fit to a four-parameter sigmoidal curve.(C)Selectivity of AA74-1 in lysates of saponin-isolated parasites. TAMRA fluorescence profiles were generated for lanes in (A) corresponding to 0, 135 and 405 nM AA74-1. The 55 and 80 kDa internalized APEH species are indicated with red asterisks. Prodrug activation and resistance esterase is indicated with a blue asterisk. For A and C, the molecular masses of markers are indicated in kDa.

To assess the selectivity of AA74-1 for APEH in saponin-isolated parasite lysate, the fluorescence profiles of the lanes corresponding to 135 and 405 nM AA74-1 in Fig. 2A were compared to that of the vehicle (DMSO) control (Fig. 2C). At both concentrations, the 80 and 55 kDa APEH species were effectively inhibited (Fig. 2C, red asterisks). A 37 kDa species was partially inhibited at both concentrations. In separate studies, we have identified this species as the “prodrug activation and resistance esterase” (R. Elahi, C. Dapper and M. Klemba, unpublished data), a serine hydrolase that is not essential for asexual replication of *P. falciparum* (31). We conclude that concentrations of AA74-1 below ∼400 nM are highly selective for the 55 and 80 kDa species of APEH in saponin-isolated parasite lysate *in vitro*.

### The anti-malarial potency of AA74-1 is enhanced over two replication cycles

To determine whether internalized APEH is required for efficient parasite replication, we examined the effect of AA74-1 on the development of a synchronized ring-stage culture. Parasite replication was assessed by measuring the fluorescence of the DNA-binding dye SYBR Green I after 48 h (the time required for one complete cycle of the 3D7 line is ∼42 h). AA74-1 was a relatively poor inhibitor of parasite growth, with incomplete inhibition of parasite replication at 10 µM and an estimated 50% effective concentration (EC_50_) greater than 1 µM (Fig. 3A, Table 1). Interestingly, however, when parasites were seeded at a lower density and allowed to proceed through two replication cycles (96 h), the efficacy of AA74-1 increased by over 10-fold, exhibiting a mean EC_50_ value of 96 ± 38 nM over three biological replicates (Fig. 3A, Table 1). Near complete inhibition of parasite replication on the second cycle was observed at an AA74-1 concentration of 310 nM, a value that is highly selective for APEH *in vitro* (Fig. 2B). In contrast to these results, parallel experiments with chloroquine yielded 48 and 96 hour EC_50_ values that were not significantly different (Table 1; Student’s two-tailed *t*-test, *p*-value = 0.93).

**Figure 3:**
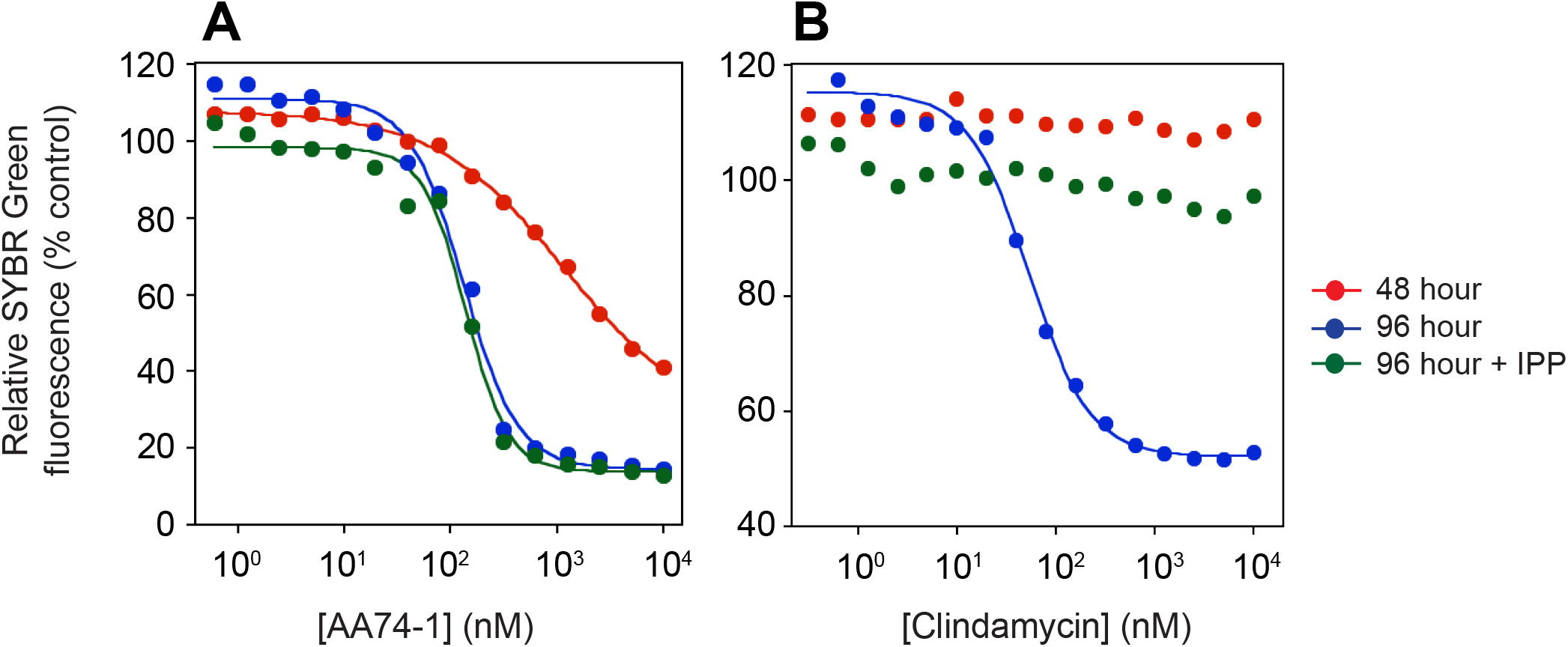
Effect of AA74-1 on parasite replication over two cycles. Concentration-response plots for parasites cultured in the presence of AA74-1 (A) or clindamycin (B) for 48 h (red circles), 96 h (blue circles) or 96 h in the presence of 200 µM IPP (green circles). At the indicated time points, parasite DNA was quantified using a SYBR Green I assay. Data are expressed as a percentage of the fluorescence value for the vehicle (DMSO) control. Inhibition curves were generated by non-linear regression fits to a four-parameter sigmoidal curve. The lower baselines are non-zero due to background fluorescence.

**Table 1:** EC_50_ values from *P. falciparum* growth inhibition assays. Values are means ± SD from three biological replicates. ND, not determined.

| Compound | 48 h EC<sub>50</sub> (nM) | 96 h EC<sub>50</sub> (nM) |  |
| --- | --- | --- | --- |
|  |  | -IPP | +IPP |
| AA74-1 | >1,000 | 96 ± 38 | 94 ± 39 |
| Clindamycin | >10,000 | 37 ± 24 | >10,000 |
| Chloroquine | 9.0 ± 2.4 | 8.5 ± 2.0 | ND |

Dramatic enhancement of anti-malarial potency during a second replication cycle is a hallmark of compounds that target the *P. falciparum* apicoplast (26, 27) and is referred to as “delayed death”. An apicoplast-based delayed death response can be reversed by supplementation of parasite culture medium with 200 µM isopentenyl pyrophosphate (IPP), which is the sole product of apicoplast metabolism that is required during the asexual growth cycle (26). To determine whether the enhanced potency of AA74-1 might be due to inhibition of an apicoplast enzyme, we conducted 96 hour growth assays in the presence and absence of 200 µM IPP. As a positive control for delayed death, parallel experiments were performed with clindamycin, an antibiotic that targets the apicoplast. Clindamycin has been shown to exhibit a profound delayed death response, which can be rescued by IPP supplementation (26, 27). While clindamycin toxicity was dramatically attenuated in the presence of IPP (Fig. 3B, Table 1), the potency of AA74-1 was unaffected (Fig. 3A, Table 1). These results strongly suggest that AA74-1 does not target an apicoplast enzyme and indicate that an alternate mechanism lies behind the enhancement of AA74-1 potency during the second replication cycle.

### Parasite-internalized APEH, but not erythrocyte APEH, is recalcitrant to AA74-1 inhibition *in vivo*

Seeking an explanation for the enhanced potency of AA74-1 during the second replication cycle, we asked whether AA74-1 is an effective inhibitor of parasite-internalized APEH *in vivo*. Because AA74-1 covalently modifies the active site serine of APEH (21), the inhibitor can be added to parasite cultures for a defined period of time and the extent of APEH inhibition *in vivo* can be assessed by TAMRA-FP labeling following inhibitor washout and saponin isolation of parasites. We determined the amount of residual internalized APEH activity following a four-hour treatment of cultured trophozoite-stage parasites with 0.1, 1 or 10 µM AA74-1. A four-hour treatment window was selected in order to minimize the potentially confounding effects of toxicity at the higher AA74-1 concentrations (Fig. 3A). Surprisingly, parasite-internalized APEH was not effectively inhibited by exogenous AA74-1 concentrations up to 10 µM (Fig. 4A). A parallel experiment with uninfected erythrocytes demonstrated robust inhibition of APEH at all exogenous AA74-1 concentrations (Fig. 4A). These results indicate that the inhibitor is able to diffuse across the erythrocyte plasma membrane but is unable to inhibit APEH within the parasite.

**Figure 4:**
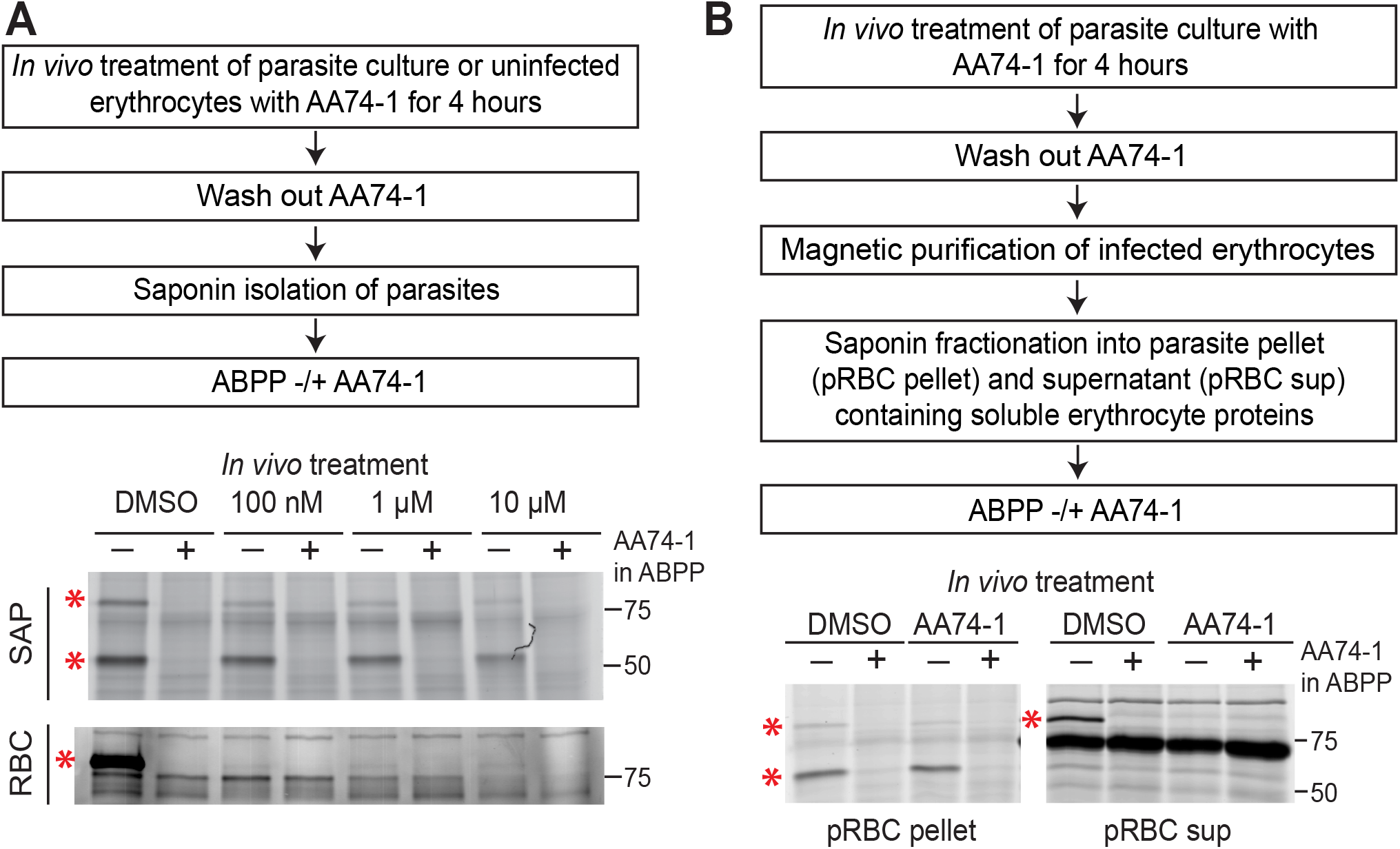
Analysis of AA74-1 inhibition of APEH *in vivo*. (A) Characterization of APEH inhibition upon AA74-1 treatment of cultured *P. falciparum*-infected or uninfected erythrocytes. Upper: Flow diagram of the experimental approach. Lower: TAMRA-FP labeling of saponin-isolated parasites (SAP) and uninfected erythrocytes (RBC) following *in vivo* treatment. (B) Characterization of *in vivo* APEH inhibition by AA74-1 in the infected erythrocyte cytosol and in saponin-isolated parasites. Upper: Flow diagram of the experimental approach. Lower: TAM-RA-FP labeling of the saponin-isolated parasite pellet (pRBC pellet) or the saponin supernatant containing soluble host erythrocyte proteins (pRBC sup) obtained from highly purified infected erythrocytes following treatment with 100 nM AA74-1. In both (A) and (B), ABPP was conducted with and without 1 µM AA74-1 (“AA74-1 in ABPP”) to identify APEH species (red asterisks). One biological replicate is shown out of two that yielded similar results. Molecular masses of markers are indicated in kDa.

To further explore this phenomenon, we conducted an experiment to determine whether there was something distinctive about *P. falciparum*-infected erythrocytes that prevented accumulation of AA74-1. Trophozoite-stage parasites were treated for four hours with 100 nM exogenous AA74-1, washed extensively, and purified on a magnetic column to > 90% parasitemia (this material is referred to as “pRBCs”). We then fractionated the pRBCs with saponin, yielding a supernatant containing host erythrocyte APEH, and a pellet containing parasite-internalized APEH. A schematic of the experimental design is shown in Fig. 4B. Once again, we observed inhibition of erythrocyte APEH but not parasitize-internalized APEH (Fig. 4B).

### Parasite-internalized APEH is active and is inhibited by AA74-1 at acidic pH

The most likely scenario for internalization of host cell APEH is through the endocytosis of large quantities of erythrocyte cytosol and delivery to the food vacuole (see Discussion). The lumen of the food vacuole is acidic with a pH of ∼ 5.5 (28, 29). To determine whether APEH could have a catalytic role at this pH, we asked whether TAMRA-FP modifies the active site serine of APEH at pH 5.5. While APEH was labeled with TAMRA-FP at pH 5.5 (Fig. 5), the extent of labeling was lower at pH 5.5 than at 7.4, which suggests a slower reaction rate at the acidic pH value. We also found that AA74-1 is capable of inhibiting APEH at pH 5.5 *in vitro* (Fig. 5); thus, an acidic pH does not by itself explain the recalcitrance of APEH to AA74-1 inhibition *in vivo*.

**Figure 5:**
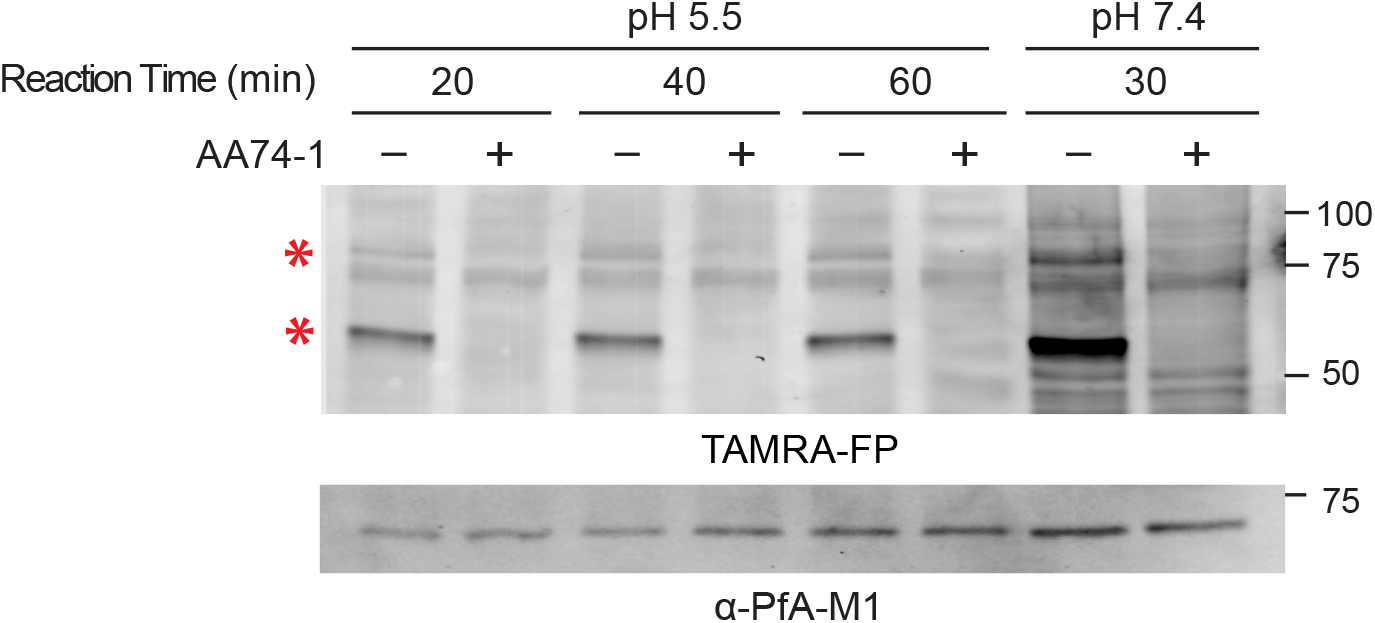
APEH activity and inhibition by AA74-1 at acidic pH. Upper panel: ABPP of saponin-isolated parasite lysates at pH 5.5 and 7.4. Times of TAMRA-FP labeling reactions are indicated. Each reaction was conducted with and without 1 µM AA74-1 to identify APEH. Lower panel: anti-PfA-M1 immunoblot to indicate relative loading levels. Red asterisks indicate the 55 and 80 kDa internalized APEH species. Molecular masses of markers are indicated in kDa.

## DISCUSSION

We present evidence that *P. falciparum* internalizes and accumulates erythrocyte APEH during its growth in the host cell. There are several reported examples of *P. falciparum* importing host cell proteins for metabolic purposes, including superoxide dismutase (30), δ-aminolevulinate dehydratase (31), and peroxiredoxin 2 (32). APEH is the first example to our knowledge of a hydrolytic enzyme accumulating in the parasite.

The most likely route for internalization of erythrocyte APEH is through the cytostomal endocytic pathway that is responsible for the uptake and delivery to the food vacuole of large quantities of erythrocyte cytosol (33–35). Up to 75% of erythrocyte hemoglobin, the dominant constituent of erythrocyte cytosol, is internalized through this pathway (36). Because cytostomal endocytosis is thought to be a non-specific process, APEH would presumably be delivered to the food vacuole along with hemoglobin and other soluble erythrocyte proteins. This model for APEH internalization is consistent with the observed proteolysis of parasite APEH to a 55 kDa species, which may be mediated by vacuolar aspartic and cysteine endopeptidases (37). Our findings are consistent with those of several previous studies demonstrating that APEH is highly resistant to proteolytic degradation and that endoproteases such as trypsin, chymotrypsin and elastase clip full-length, 80 kDa APEH into ∼55 kDa and ∼25 kDa fragments *in vitro* (38–41). Interestingly, these studies have established that this proteolytic treatment neither disrupts the homotetrameric structure of APEH nor reduces its activity (38, 39), thus providing a plausible explanation for the stability of APEH in the proteolytic environment of the food vacuole. The shift in size of APEH in saponin-isolated parasite lysates indicates that it does not originate from contamination by host erythrocyte cytosol or from interaction with the outer leaflet of the parasite plasma membrane. We have tried to confirm a food vacuole location for APEH through indirect immunofluorescence using anti-APEH antibodies; however, although we have tested a wide range of fixation conditions and numerous antibodies, we have not found conditions that are suitable for detection of APEH in infected erythrocytes.

To investigate the possibility of a physiological role for internalized APEH, we employed the APEH-selective inhibitor AA74-1 that was discovered in a library of triazole urea compound by Cravatt and colleagues (21). The exquisite selectivity of this covalent inhibitor for APEH has been demonstrated *in vitro* using mouse T cell lysates and *in situ* using cultured T cells (21). Furthermore, upon treatment of mouse T cells with AA74-1, the N-terminal acetylation state of ∼25 endogenous proteins was altered, which indicates that AA74-1 is able to engage its target *in situ* (21). Thus, we considered AA74-1 to be an appropriate tool for the interrogation of APEH function in *P. falciparum*. To validate the use of AA74-1 in the context of *P. falciparum*-infected erythrocytes, we demonstrated by competitive ABPP that the inhibitor is highly selective for parasite-internalized APEH in lysates of saponin-isolated parasites at concentrations below ∼400 nM. It is also selective for APEH in uninfected erythrocytes (Fig. S1).

When AA74-1 was added to synchronized ring-stage parasites, inhibition of growth over the first replication cycle required concentrations that were much higher than those needed to inhibit APEH *in vitro*. However, a dramatic enhancement of potency was observed if the growth inhibition experiment was continued for a second replication cycle, *i.e.* 96 hours. Notably, the EC_50_ for growth inhibition after the second cycle was only 12-fold higher than the IC_50_ observed for inhibition of APEH in competitive ABPP *in vitro*. Furthermore, the AA74-1 concentrations that yielded efficient inhibition of growth replication were within the concentration range found to be highly selective by competitive ABPP (*i.e.*, below 400 nM). Taken together, the evidence strongly suggests that the growth defect observed during the second cycle is due to the selective targeting of APEH.

Enhancement of drug potency during a second replication cycle is commonly observed with inhibitors that target the parasite apicoplast (42). Our experiments with IPP supplementation, however, revealed that internalized APEH is not acting in the context of the apicoplast, nor is AA74-1 cross-inhibiting an apicoplast enzyme. Rather, it became clear from *in vivo* experiments (*i.e.*, upon addition of AA74-1 to the medium of cultured parasites) that the parasite-internalized APEH species were not inhibited by AA74-1 under these conditions. This outcome was not due to its inability to penetrate the host cell: the erythrocyte pool of APEH was quantitatively inhibited by AA74-1, both in uninfected erythrocytes and in the erythrocyte cytosol of infected cells. The reason for the recalcitrance of parasite-internalized APEH is not entirely clear. If APEH resides in the food vacuole, it will experience an acidic pH; however, we have demonstrated that the 55 kDa APEH species reacts with AA74-1 at pH 5.5 *in vitro*. It is possible that AA74-1 is not able to access the lumen of the food vacuole, or that it is inactivated inside the parasite by a hydrolytic enzyme. Whatever the reason, it is plausible that any parasite-internalized APEH that is present at the initiation of a growth inhibition experiment remains active throughout the replication cycle. The enhancement of potency during the following replication cycle could derive from the fact that the parasites invade erythrocytes containing AA74-1-inactivated APEH and therefore internalize inactive enzyme.

Our findings lead us to the intriguing idea that *P. falciparum* has adapted to use internalized APEH for a crucial metabolic function. The most apparent role for APEH, if as expected it resides in the food vacuole, is in catalyzing the hydrolysis of acetylated amino acids from the N-termini of peptides generated through the catabolism of endocytosed erythrocyte proteins. The two most abundant cytosolic proteins in the erythrocyte are hemoglobin (composed of α– and β–globin in the adult) and carbonic anhydrase-1, which are present at 97% and 1% of total protein, respectively (43). While α– and β–globin have not traditionally been thought to have acetylated N-termini, a proteome-wide analysis of erythrocyte proteins that explicitly addressed the acetylation status of N-termini found that the dominant species of both α– and β–globin are N-acetylated at a frequency of about 20% (44). Carbonic anhydrase-1 is known to possess an acetylated N-terminus (45). Furthermore, proteomic studies have identified over 1,500 soluble erythrocyte proteins (46, 47), around 53% of which are N-terminally acetylated (44). Given the large quantities of erythrocyte cytosol that are digested in the food vacuole, a mechanism is likely needed for efficient removal of N-acetylated amino acids from peptides generated by endoproteolytic hydrolysis, as N-blocked peptides are expected to be poor substrates for the vacuolar exopeptidase dipeptidyl aminopeptidase 1 and the M1- and M24-family aminopeptidases PfA-M1 and PfAPP (48, 49). Although the pH optimum of human erythrocyte APEH is reported to be close to neutral pH (13, 50), TAMRA-FP labeling at pH 5.5 strongly suggests that parasite-internalized APEH retains catalytic activity, albeit diminished, at the acidic pH of the food vacuole. The co-internalization of host cell APEH could provide the parasite with an elegant solution for the need to catabolize N-acetylated peptides. Our findings lay the groundwork for further studies (such as metabolomic approaches) that could provide a direct test of this hypothesis.

## MATERIALS AND METHODS

### Reagents

The TAMRA-fluorophosphonate activity-based probe was obtained from ThermoFisher. *N*-(*trans*-epoxysuccinyl)-*L*-leucine 4-guanidinobutylamide (E-64), AA74-1, clindamycin, chloroquine and isopentenyl pyrophosphate trilithium salt were purchased from Sigma. Pepstatin A was purchased from MP biomedicals.

### Parasite culture

*P. falciparum* 3D7 was cultured in human O^+^ erythrocytes (Interstate Blood Bank, Memphis, TN) at 2% hematocrit in RPMI 1640 medium supplemented with 27 mM sodium bicarbonate, 11 mM glucose, 0.37 mM hypoxanthine, 10 µg/mL gentamicin, and 5 g/L Albumax I (Invitrogen). Unless otherwise indicated, cultures were incubated at 37 °C in a 5% CO_2_ incubator. Cultures were synchronized by treatment with 5% (w/v) sorbitol (51).

### Preparation of parasite lysate

Synchronized cultures of maturing parasites (32-40 h post-invasion, hpi) were separated from soluble erythrocyte and parasitophorous vacuole proteins by treatment with 0.03% (w/v) saponin in cold Dulbecco’s phosphate-buffered saline (PBS, pH 7.4) for 10 minutes on ice. Saponin-isolated parasites were recovered by centrifugation at 1940 × *g* at 4 °C for 10 minutes and were washed three times with cold PBS. The yield of parasites was determined by counting on a hemocytometer. Parasites were suspended to a density of 5 × 10^8^ parasites/mL in cold PBS containing the protease inhibitors pepstatin A (5 µM) and E-64 (10 µM). The parasite suspension was subjected to three rounds of sonication at 30% power for 10 seconds. After centrifugation at 17,000 × *g* to pellet cellular debris, aliquots of clarified lysates were snap frozen in liquid N_2_ and stored at – 80 °C.

### Preparation of uninfected erythrocyte lysate

Uninfected erythrocytes were washed thrice in cold PBS, counted on a hemocytometer and resuspended in cold PBS containing 5 µM pepstatin and 10 µM E-64 to a density to 5 × 10^8^ cells/mL. Lysates of resuspended erythrocytes were prepared and stored as described above for saponin-isolate parasites.

### Activity-based protein profiling

TAMRA-FP labeling reactions were conducted with 19.8 µL of parasite or erythrocyte lysate, which corresponds to ∼ 10^7^ cells/reaction. To start the reaction, 0.2 µL of 100 µM TAMRA-FP was added, giving a final concentration of 1 µM. Reactions were incubated at 30 °C for 30 minutes and then stopped by the addition of one volume of 2x reducing SDS-PAGE loading buffer and incubation at 95 °C for 5 minutes. For competitive ABPP experiments, inhibitor or vehicle (DMSO) was added and reactions were incubated for 20 minutes at 30 °C prior to the addition of TAMRA-FP. Labeled proteins were resolved on 8.5% or 10% reducing SDS-polyacrylamide gels. In-gel TAMRA fluorescence was recorded on a Typhoon Trio flatbed scanner (GE Healthcare Life Sciences, Piscataway, NJ). Fluorescence profiles and peak volumes of labeled proteins were obtained using ImageQuant TL v2005 (GE Healthcare Life Sciences, Piscataway, NJ). For calculation of AA74-1 IC_50_ values (Fig. 2), the peak volume for the 55 kDa APEH species in saponin-isolated parasites was normalized to that of a ∼160 kDa species (Fig. 2A, black asterisk) that was not inhibited by AA74-1 at any concentration. Normalization of erythrocyte APEH peak volumes was conducted in a similar manner (see Fig. S2). IC_50_ values were calculated by nonlinear regression fitting of the data to a four-parameter sigmoidal curve using KaleidaGraph 4.5 (Synergy software, Reading, PA).

### Immunoblotting

Competitive ABPP on crude lysates from ∼ 10^7^ saponin-isolated parasites or uninfected erythrocytes was performed as described above. Following in-gel fluorescence scanning, proteins were transferred to a nitrocellulose membrane which was blocked with 2% bovine serum albumin in Tris-buffered saline containing 0.1% Tween 20 (TBST/BSA) for one hour at room temperature. The membrane was then incubated with primary antibody diluted in TBST/BSA for one hour followed by a one hour incubation with horseradish peroxidase-conjugated anti-rabbit secondary antibody (1:10,000, GE Healthcare life sciences, Piscataway, NJ). Primary antibodies used were: affinity purified anti-APEH rabbit polyclonal (IgG) raised against an APEH fragment consisting of amino acids 381-732 (product # 14758-1-AP, Proteintech, Rosemont, IL; 0.26 µg/mL) and affinity purified anti-PfA-M1 (0.13 µg/mL; (52)). Chemiluminescent signal was developed with ECL Plus (GE Healthcare life sciences, Piscataway, NJ) and recorded on x-ray film, which was digitized by scanning, or imaged on a ChemiDoc MP system (Bio-Rad laboratories, Hercules, CA). Image contrast for the TAMRA-FP fluorescence and chemiluminescent signal was adjusted with Adobe Photoshop CS2 (Adobe, Inc., San Jose, CA).

### *P. falciparum* growth inhibition assays

Synchronized ring-stage cultures were seeded at 3% parasitemia (48 h assay) or 0.6% parasitemia (96 h assay) and 1% hematocrit in a 96-well flat bottom plate. Inhibitors were added from 1000x stock solutions in DMSO to generate two-fold concentration series of AA74-1 (0.61 nM – 10 µM), chloroquine (1.9 – 250 nM) or clindamycin (0.3 nM – 10 µM). After 48 or 96 h incubation at 37 °C under reduced oxygen conditions (5% O_2_, 5% CO_2_, and 90% N_2_), parasite growth was determined using a SYBR Green I DNA quantitation assay as previously described (53). Values from samples containing 0.1% DMSO were used to calculate relative SYBR Green fluorescence. Each assay was performed with two technical replicates, which were averaged to generate a single biological replicate. EC_50_ values were calculated by nonlinear regression fitting of the data to a four-parameter sigmoidal curve using KaleidaGraph 4.5 (Synergy software, Reading, PA). Means and standard deviations from three biological replicates are reported in Table 1. For experiments with 200 µM IPP supplementation, single technical replicates were conducted for each biological replicate.

### AA74-1 inhibition of APEH *in vivo*

Trophozoite-stage parasites (30-38 hpi) or uninfected erythrocytes were incubated in culture medium supplemented with AA74-1 (100 nM, 1 µM, or 10 µM) or with 0.1% DMSO for four hours at 37 °C under reduced oxygen conditions with gentle mixing on an orbital rotator. Cultures were washed four times in cold RPMI to remove exogenous AA74-1. Parasites were then isolated with saponin as described above. Saponin-isolated parasites and uninfected erythrocytes were counted with a hemocytometer, resuspended in cold PBS at a cell density of 5 × 10^8^ per mL, and stored at –80 °C. Samples were assayed directly by competitive ABPP.

To investigate the inhibition of APEH in the host and parasite compartments of infected erythrocytes, a synchronized trophozoite culture was treated with AA74-1 (100 nM) or with 0.1% DMSO for four hours at 37 °C under reduced oxygen conditions. The culture was washed four times in cold RPMI to remove AA74-1 and then resuspended in RPMI media. Parasitized erythrocytes were purified from the culture on a MACS magnetic LD column (Miltenyi Biotech, Gaithersburg, MD) following the manufacturer’s instructions. Enriched parasitized RBCs (pRBCs) were subjected to saponin treatment as described in the section “Preparation of parasite lysate”. Saponin-isolated parasites (pRBC pellet) and the supernatant containing soluble host erythrocyte proteins (pRBC sup) were collected and stored at –80 °C. Samples were used directly for competitive ABPP.

### Activity and inhibition of APEH at acidic pH

Saponin-isolated parasites were split into two aliquots and resuspended to a density of 5 × 10^8^ parasites/mL in either PBS pH 7.4 or 100 mM sodium 2-(*N*-morpholino)ethanesulfonate (MES) pH 5.5, both of which included 5 µM pepstatin A and 10 µM E-64. Crude lysates were prepared as described in “Preparation of parasite lysate”. APEH activity and inhibition by AA74-1 were analyzed as described in “Activity-based protein profiling”, with the modification that TAMRA-FP incubation times up to 60 minutes were employed for reactions conducted at pH 5.5. After fluorescence scanning, proteins were immediately transferred to nitrocellulose and relative loading levels were assessed by immunoblotting with an antibody against the *P. falciparum* aminopeptidase PfA-M1 (see “Immunoblotting”).

## ACKNOWLEDGEMENTS

This work was supported by National Institute of Allergy and Infectious Diseases grant AI133136. The funding agency had no role in study design, data collection and interpretation, or the decision to submit the work for publication. R.E. and M.K. designed the experiments, R.E. and C.D. conducted the experiments, and R.E. and M.K. prepared the manuscript.

## FIGURE LEGENDS

**Figure S1:**
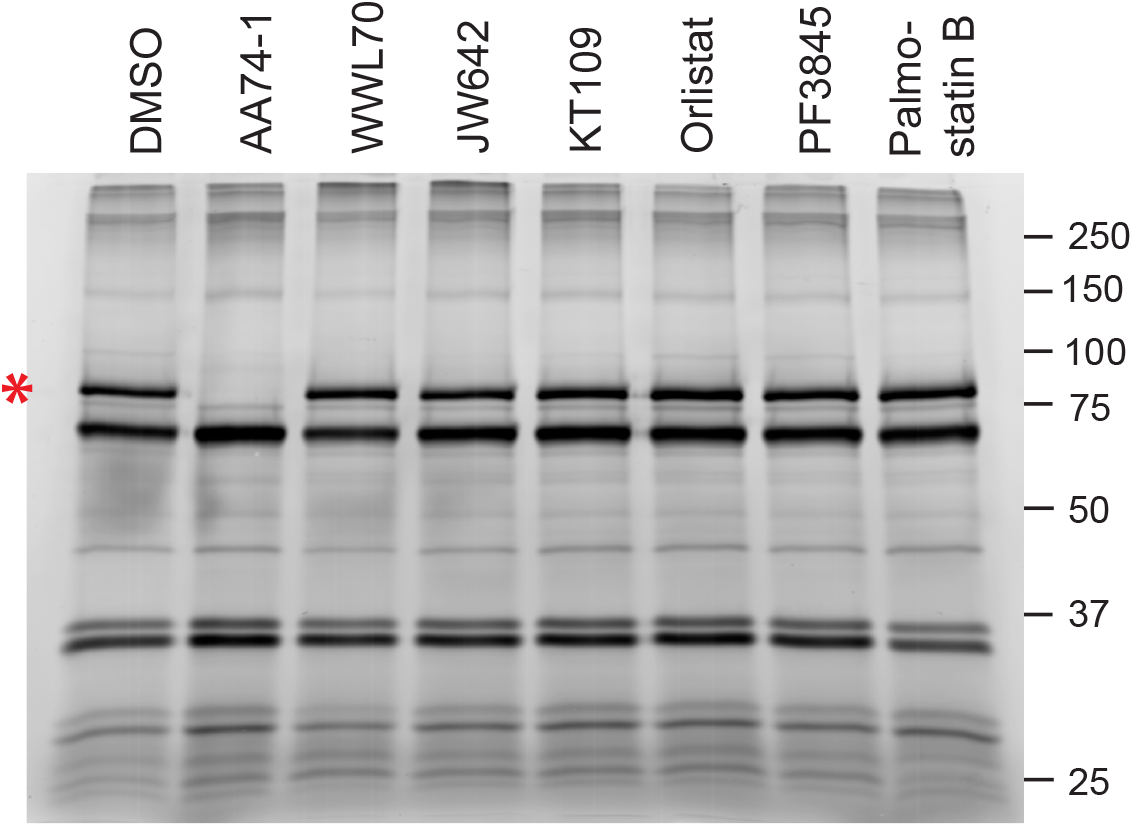
Competitive ABPP of uninfected erythrocyte lysate. APEH is indicated with a red asterisk. Inhibitor structures and selectivities are given in Table S1. Molecular masses of markers are indicated in kDa.

**Figure S2:**
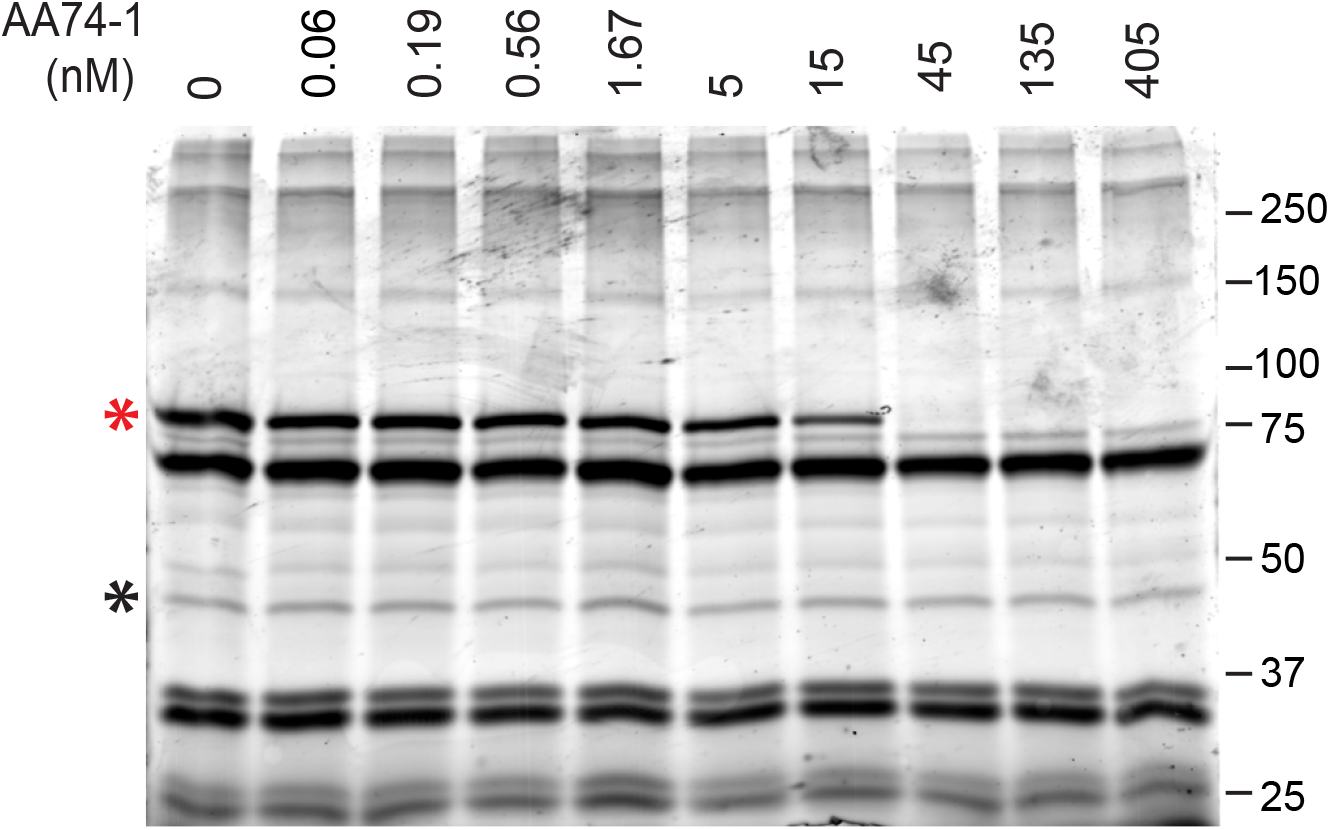
*In vitro* potency and selectivity of AA74-1 for erythrocyte APEH. The full gel corresponding to the segment in Fig. 2A is shown. APEH is indicated with a red asterisk and the species used for peak volume normalization is indicated with a black asterisk. Molecular masses of markers are indicated in kDa.

---

**Table S1:**
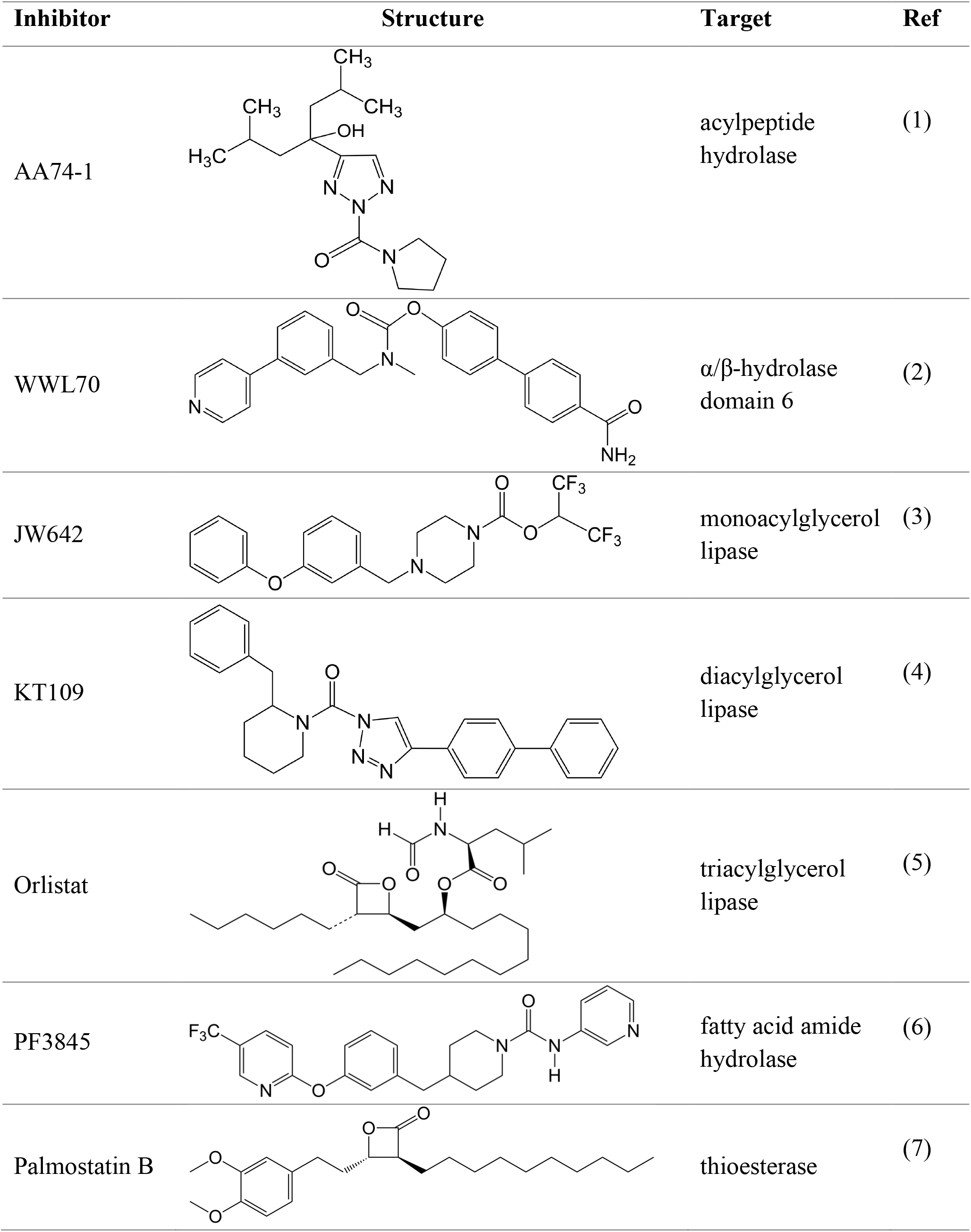
Serine hydrolase inhibitors used in this study for competitive activity-based protein profiling.

| Inhibitor | Structure | Target | Ref |
| --- | --- | --- | --- |
| AA74-1 |  | acylpeptide hydrolase | (1) |
| WWL70 | | $\alpha/\beta$-hydrolase domain 6 | (2) |
| JW642 |  | monoacylglycerol lipase | (3) |
| KT109 |  | diacylglycerol lipase | (4) |
| Orlistat |  | triacylglycerol lipase | (5) |
| PF3845 |  | fatty acid amide hydrolase | (6) |
| Palmostatin B |  | thioesterase | (7) |

